# LiGIoNs: A Computational Method for the Detection and Classification of Ligand-Gated Ion Channels

**DOI:** 10.1101/833350

**Authors:** Avgi E. Apostolakou, Katerina C. Nastou, Georgios N. Petichakis, Zoi I. Litou, Vassiliki A. Iconomidou

## Abstract

Ligand-Gated Ion Channels (LGICs) are one of the largest groups of transmembrane proteins. Due to their major role in synaptic transmission, both in the nervous system and the somatic neuromuscular junction, LGICs present attractive therapeutic targets. During the last few years several computational methods for the detection of LGICs have been developed. These methods are based on machine learning approaches utilizing features extracted solely from amino acid composition. Here we report the development of LiGIoNs, a profile Hidden Markov Model (pHMM) method for the prediction and ligand-based classification of LGICs. The method consists of a library of 10 pHMMs, one per LGIC subfamily, built from the alignment of representative LGIC sequences. In addition, 14 Pfam pHMMs are used to further annotate and classify unknown protein sequences into one of the 10 LGIC subfamilies. Evaluation of the method showed that it outperforms existent methods in the detection of LGICs. On top of that, LiGIoNs is the only currently available method that classifies LGICs into subfamilies.

The method is available online at http://bioinformatics.biol.uoa.gr/ligions/.

## 1. INTRODUCTION

Most living cells exhibit a membrane potential due to the membrane acting as a barrier for charged particles [1]. To overcome this obstacle, cells use specialized transmembrane proteins – known as ion channels [2] – that control the flow of ions in and out of the cell. These proteins are highly selective and can discriminate both between anions and cations as well as between monovalent and divalent ions [3]. Their gating is typically a result of either change to the membrane potential or binding of specific ligands. Thus, channels are typically classified in two main classes, according to their gating trigger, Voltage-Gated Ion Channels (VGICs) and Ligand-Gated Ion Channels (LGICs) [4]. Both classes are extremely diverse and composed of numerous members further classified into various families and subfamilies.

Most LGICs consist of multiple subunits forming heteropolymers; these subunits are encoded by many genes. The variety of possible subunit combinations within each subfamily of LGICs leads to a wide range of receptors with different pharmacological and biophysical properties and diverse expression patterns both within the nervous system and in other tissues [5]. Thus, LGICs have emerged as complex yet attractive targets for the development of new therapeutic agents [6]. LGICs comprise the anion-selective GABA_A_ receptors [7] and glycine receptors [8], and the cation-selective nicotinic acetylcholine receptors [9,10], 5-HT_3_ receptors [11], ionotropic glutamate receptors [12], IP_3_ receptors [13], P2X receptors [14], epithelial sodium channels [15], acid-sensing (proton-gated) ion channels [16] and zinc-activated channels [17]. The nicotinic acetylcholine, 5-HT_3_, GABA_A_ and glycine receptors (and an additional member, the zinc-activated channels) form the family of Cys-loop receptors [18]. The distinct structural characteristics of each LGIC subfamily are shown in **Table 1**.

**Table 1.** Characteristics of the 10 LGIC subfamilies. The number of subunits, transmembrane segments per subunit and average length of each subunit in amino acid residues is shown for each subfamily. Members of the Cys-loop family are marked with *italics* and underline in the table.

| *LGIC subfamily* | *Subunit Count* | *Transmembrane (TM) segments per subunit* | *Subunit Length* |
| --- | --- | --- | --- |
| Epithelial Sodium Channels (ENaCs) | 3 | 2 | ~650 |
| P2X Receptors | 3 | 2 | ~435 |
| Acid-sensing Ion Channels (ASICs) | 3 | 2 | ~550 |
| <u>*Ionotropic Glutamate Receptors*</u> | 4 | 3 | ~950 |
| <u>*Nicotinic Acetylcholine Receptors*</u> | 5 | 4 | ~500 |
| <u>*5-HT<sub>3</sub> Receptors*</u> | 5 | 4 | ~460 |
| <u>*GABA<sub>A</sub> Receptors*</u> | 5 | 4 | ~480 |
| <u>*Glycine Receptors*</u> | 5 | 4 | ~470 |
| <u>*Zinc-activated Channels (ZACs)*</u> | 5 | 4 | ~410 |
| IP<sub>3</sub> Receptors | 4 | 6 | ~2750 |

Considering the importance of ion channels for normal cellular function and their role as drug targets [19,20], several prediction methods have been developed for the identification of ion channels based on their amino acid sequence [21–29]. In a recent survey [30] ion channel predictors were compared and evaluated and only two of those, IonChanPred 2.0 [27] and PSIONplus [26], were both available for use and capable of classifying ion channels into LGICs. IonChanPred 2.0 is built on an SVM-based model that employs pseudo-dipeptide composition and physicochemical properties. PSIONplus combines SVM and PSI-BLAST and relies on multiple features; it performed better than IonChanPred 2.0, but still modestly so. Both methods were designed to work in parallel in order to detect ion channels, then classify them into LGICs or VGICs and in the case of VGICs to further classify them into four subtypes according to ion selectivity. Since the publication of the survey a new predictor by Han *et al*. [28] came out, that is however not available, and PSIONplus^m^ [29], built on PSIONplus, that addressed some issues pointed out by the survey, including now classifying LGICs into four sub-types, same as VGICs. However, there is no available method that classifies LGICs into subfamilies.

In order to fill this gap, we decided to design and develop LiGIoNs, a sequence-based predictor, that identifies LGICs in proteomes with the use of profile Hidden Markov Models (pHMMs). Each pHMM describes a different LGIC subfamily and can be used to detect novel members. As such, LiGIoNs is the first method that in addition to detecting LGICs, performs classification of LGICs into the 10 known ligand-based sub-families (**Table 1**). We have also developed a web server to host the method, available at http://bioinformatics.biol.uoa.gr/ligions.

## 2. METHODS

The LiGIoNs method consists of two parts: the prediction part (detection and classification) and the annotation part. For the prediction part of the method, a dataset of LGICs was collected for each subfamily following the IUPHAR classification scheme [31] already presented in **Table 1**. For the training set, proteins were originally collected from the IUPHAR database [31]. However, since IUPHAR contains data only for human, mouse and rat proteins, the exclusive use of this source would greatly limit the diversity of the training dataset. For this reason, we isolated all UniProt [32] entries with evidence at protein or transcript level (an exception was made for ZACs) that belong to an LGIC subfamily (UniProt queries found in **Supplementary Table 1**). These proteins were cross-checked with the IUPHAR database and only two proteins in IUPHAR were excluded due to their existence evidence in UniProt. The final dataset of protein sequences used for the LiGIoNs training dataset is shown in **Supplementary Table 1**. The CD-HIT clustering algorithm [33,34] was used to reduce sequence similarity to 40% within each subfamily. The remaining members in each subfamily were aligned using ClustalO [35], resulting in 10 multiple sequence alignments (MSA). HMMER 3.3.2 [36] was then used to construct the 10 respective pHMMs corresponding to the MSA of each LGIC subfamily, which all together constitute the LGICslib pHMM library.

The following algorithm is used to characterize an unknown protein sequence as an LGIC. Initially, the unknown protein sequence is scanned against LGICslib. This is followed by recording the number of pHMMs that align with the unknown protein sequence. In order to characterize an unknown protein sequence as a member of a specific LGIC subfamily the alignment score must be higher than the threshold that is set for that pHMM. The thresholds for each pHMM were set manually in order to maximize sensitivity and specificity, based on the protocol introduced by Ioannidou *et al*. [37].

For the annotation part of LiGIoNs, all characteristic pHMMs for the LGICs were identified and isolated from the Pfam protein family database [38]. This procedure preceded the construction of the LGICslib pHMMs, in an initial attempt to identify existing pHMMs that could be used to uniquely describe LGIC subfamilies. However, this was not possible using data extracted exclusively from Pfam, as there is no combination of pHMMs deposited in the database that allows the successful classification of LGICs into subfamilies. Nevertheless, pHMMs from Pfam (**Table 2**) in combination with those in LGICslib, allowed both an additional validation of the results obtained using LiGIoNs in proteomes and the creation of the annotation part of the method.

**Table 2.**
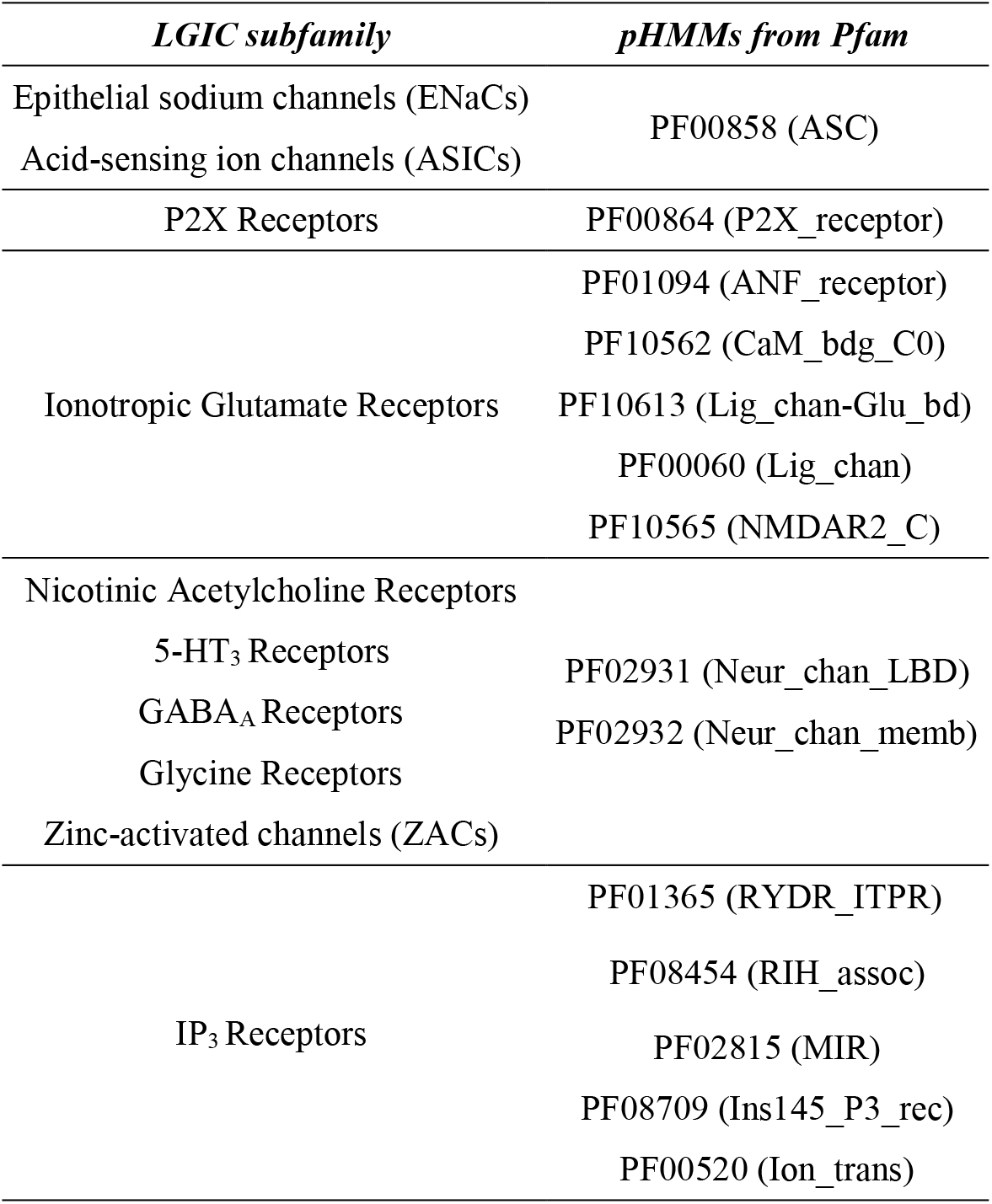
Correlation between pHMMs deposited in Pfam and LGIC subfamilies.

The pHMMs that are characteristic of LGICs were collected from the Pfam cross-references provided in all UniProt entries of the LGICs training set (**Supplementary Table 1**). As a result, 14 pHMMs were isolated from Pfam, which are presented in **Table 2** and were used to create a second library, named PfamLGICslib. HMMER is used again to scan sequences – previously characterized as LGICs by LiGIoNs – against PfamLGICslib, and sequences that have a “hit” from pHMMs in the Pfam library are annotated with these domains.

A jackknife cross-validation experiment was conducted to assess the performance of LiGIoNs, by evaluating the performance of pHMMs in correctly classifying LGICs belonging to different subfamilies. For each LGIC subfamily, one sequence was removed from the multiple sequence alignment (MSA) of the pHMM’s seed set, and a new pHMM was constructed from the remaining sequences of the new MSA. Then we measured the newly created pHMM’s ability to correctly classify the removed sequence and the sequences from three negative datasets. The three negative datasets consisted of (1), a set of all LGICs of the other subfamilies, (2) a set of non-LGIC transmembrane proteins and (3) a set of globular proteins. The set of all LGICs of other families includes the protein members of each subfamily after the sequence similarity reduction. The negative test set of non-LGIC transmembrane proteins was isolated by searching UniProt/SwissProt for transmembrane proteins with many transmembrane segments (subcellular location: “Multi-pass membrane protein”) not containing the keyword “Ligand Gated Ion Channel” in the entry’s text file. This search returned 52581 entries. An additional dataset of globular proteins was collected from UniProt/SwissProt (excluding subcellular location: “Multi-pass membrane protein” or “Single-pass membrane protein”) of which 1000 were selected at random. Both datasets were subjected to sequence similarity reduction using CD-HIT to below 40% similarity resulting in a representative set of 1508 transmembrane proteins and a set of 953 globular proteins (**Supplementary Table 2**).

In addition to the above evaluation, the method was compared to the IonChanPred 2.0 method to test its overall performance. The negative dataset used by IonChanPred 2.0 [27] for performance evaluation, comprised of 300 non-LGIC transmembrane proteins, was used to compare the two methods, in addition to the aforementioned datasets. A comparison to the PSIONPlus^m^ method [29] was not feasible, as the online tool is prohibitively slow. It should also be mentioned that none of the above methods are capable of classifying LGICs into subfamilies.

True/false positives (TP, FP) and true/false negatives (TN, FN) were counted on a per protein basis. Specifically, for the jackknife cross-validation a protein in the negative dataset was counted as a FP if it is incorrectly classified as LGIC in at least one iteration and as a TN if it is always correctly classified as non-LGIC. For the prediction performance of LiGIoNs three measures were used, namely Recall, Precision and F-score. These measures are best suited for binary classification and are not affected by the number of TN [39], necessary for the imbalanced datasets used in this study.

Recall, or Sensitivity is:

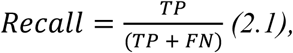

Precision is:

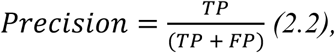

and F-score is:

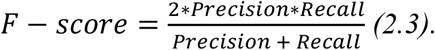

Moreover, LiGIoNs was applied to all reference eukaryotic, bacterial and archaean proteomes (**Supplementary Table 3**) retrieved from UniProt (release: 2021_02) in order to further assess the method’s ability to detect LGICs in proteomes. An additional measure was used for the proteome analysis, the False Positive Rate (FPR), to assess the number of proteins expected to be predicted as LGICs by chance given the size of each respective proteome.

FPR is calculated as:

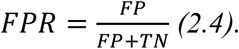

## 3. RESULTS AND DISCUSSION

### 3.1. LiGIoNs algorithm

The prediction part of LiGIoNs consists of a library of 10 pHMMs (LGICslib), created from the MSAs of the proteins belonging to the 10 LGIC subfamilies (Table 3), after sequence similarity reduction. For a protein to be characterized as a subunit of a specific LGIC subfamily, the corresponding pHMMs in LGICslib must score higher than the profile’s threshold. If multiple subfamilies meet the aforementioned condition, the one with the highest overall score is chosen to characterize the unknown sequence, but all hits are made available to the user for further consideration. For the annotation level, a library of 14 pHMMs containing characteristic LGIC domains recorded in Pfam was created (PfamLGICslib, **Table 2**). This library is scanned in positive cases only – i.e. an unknown sequence is characterized as an LGIC in the previous step – in order to provide the user with more information regarding the protein being studied. The flowchart in **Figure 1** depicts in detail how the LiGIoNs method works.

**Table 3.**
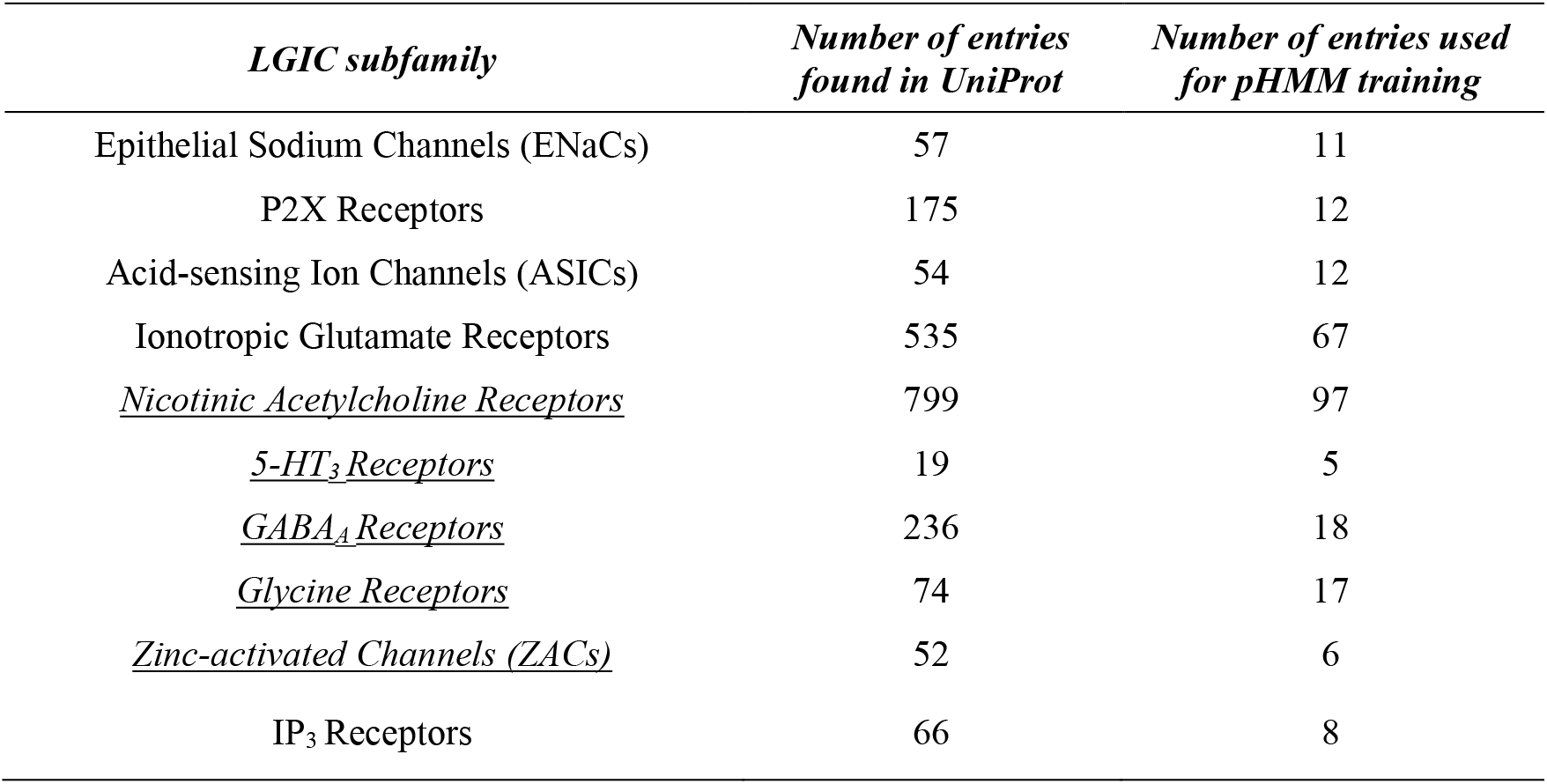
Dataset of LGICs per subfamily. LGICs were collected from UniProt and the training datasets were created after a reduction of sequence similarity at 40%. (More details in **Supplementary Table 1**)

| *LGIC subfamily* | *Number of entries found in UniProt* | *Number of entries used for pHMM training* |
| --- | --- | --- |
| Epithelial Sodium Channels (ENaCs) | 57 | 11 |
| P2X Receptors | 175 | 12 |
| Acid-sensing Ion Channels (ASICs) | 54 | 12 |
| Ionotropic Glutamate Receptors | 535 | 67 |
| <u>Nicotinic Acetylcholine Receptors</u> | 799 | 97 |
| <u>5-HT<sub>3</sub> Receptors</u> | 19 | 5 |
| <u>GABA<sub>A</sub> Receptors</u> | 236 | 18 |
| <u>Glycine Receptors</u> | 74 | 17 |
| <u>Zinc-activated Channels (ZACs)</u> | 52 | 6 |
| IP<sub>3</sub> Receptors | 66 | 8 |

**Figure 1.**
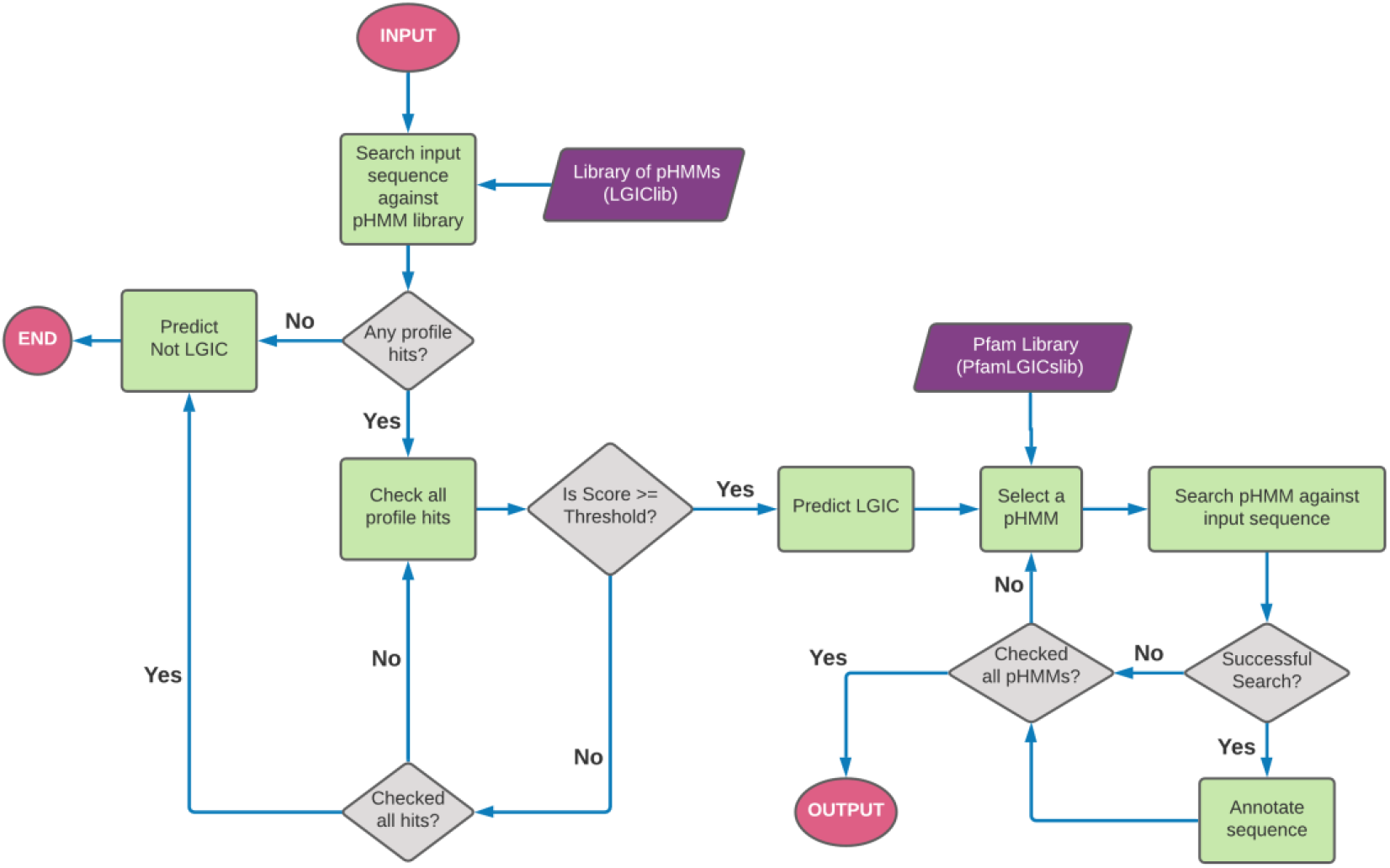
Flowchart of the LiGIoNs algorithm. A FASTA formatted sequence is used as input. The input sequence is used to search against the pHMM library LGICslib. If the search is successful, the profile hits are checked for the highest scoring hit that has a score above the corresponding profile threshold. If these conditions are not met, then the sequence is characterized as a non-LGIC and the program exits. On the other hand, if the conditions are met the sequence is characterized as LGIC and is then scanned against the library of Pfam profiles (PfamLGICslib). After all profiles have been searched against the sequence, the annotation procedure is over, and the results are presented to the user.

### 3.2. User interface and website features

A web interface has been created for LiGIoNs and the method is publicly available through http://bioinformatics.biol.uoa.gr/ligions. Via the “Home” page, the user can access the query submission page (“Run”) and the “Manual” page. Query submission can be performed using a single or a set of protein sequences in the textbox provided, or by uploading a file with FASTA formatted sequences (up to 3000 sequences). After a successful query submission, users are transferred to the results page where they can gather information about their submission and download a file with the results. In the same page users can download the results in a text file and perform a new query.

At the “Results” page, all results are presented in a table format, where each line corresponds to a single protein and each column contains the basic characteristics of the proteins and the prediction of LiGIoNs (LGIC or no LGIC). More details about each protein are provided, as shown in **Figure 2**.

**Figure 2.**
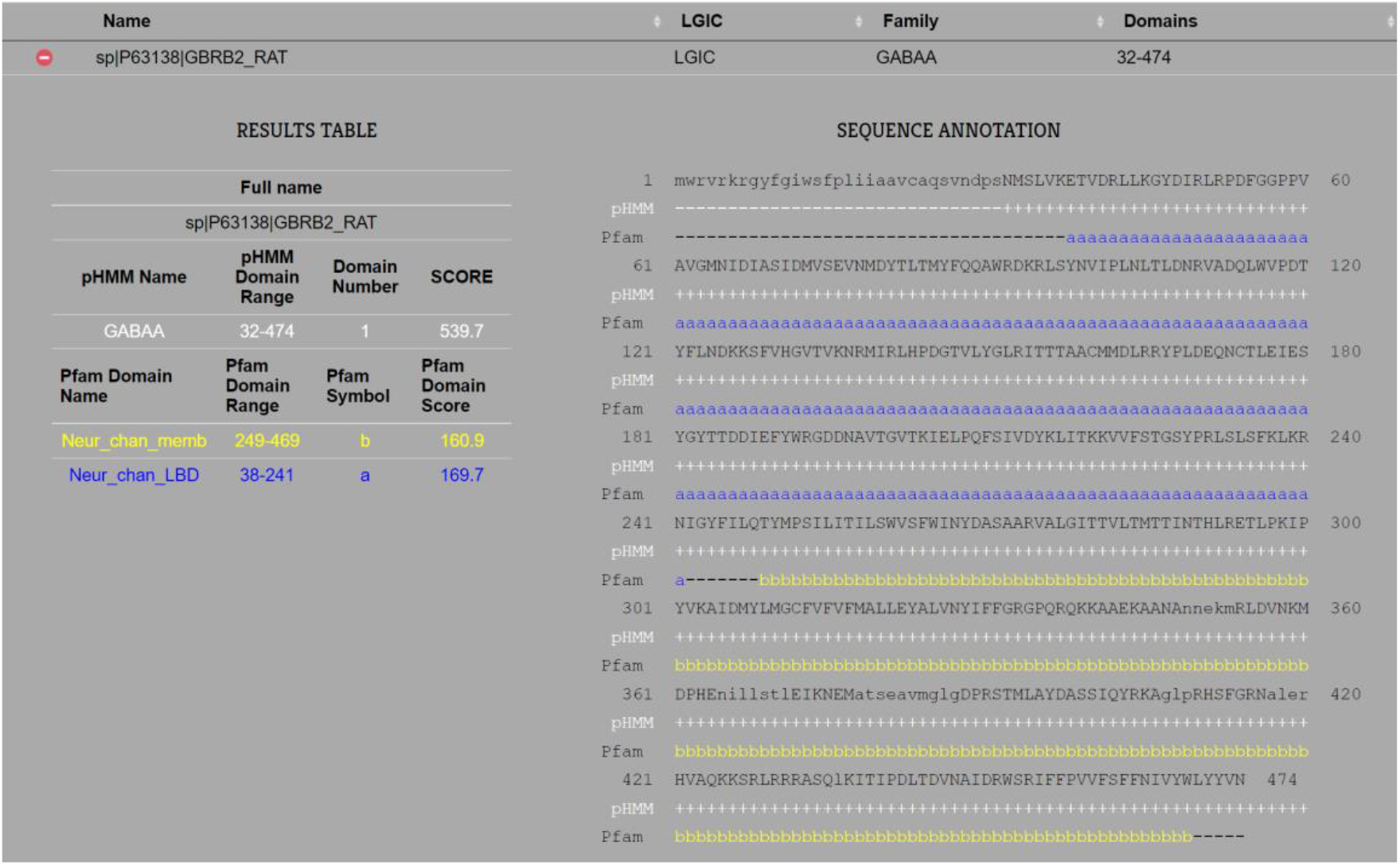
Webpage results after a successful run of LiGIoNs against an unknown protein sequence. The sequence of the GABA_A_ receptor from *Rattus norvegicus* (Rat) (UniProtAC: P63138) is used in this example. The pHMM of LGICslib that corresponds to GABA_A_ receptors, as well as two pHMMs from PfamLGICslib, characteristic of LGICs that belong to the Cys-loop family (Table 2) are detected in the sequence. The first line under the sequence displays the positions where the input sequence and the GABA_A_ pHMM are aligned (+ symbol). The second line under the sequence shows the positions where the different Pfam domains have been detected in the sequence, using a different color and character for each one of them. Positions where multiple domains have been detected, are marked with asterisks (*).

The results text file contains a protein identifier, the protein subfamily that the protein belongs to – if it is a positive hit, as well as other subfamily hits and their scores, the position and score of each domain, Pfam domain(s) present in the protein, the protein sequence and the alignment annotation from HMMER. Users are provided with a JobID for each submission, which can be used for up to two weeks to retrieve results after a prediction has been performed. LiGIoNs is fast, since for a query length the size of the human proteome the method produces results in approximately two hours, which makes it well-suited for proteomic scale applications. Lastly, in addition to the web interface all scripts and associated files needed to run LiGIoNs or recreate the pHMMs are available here https://github.com/DawnBio/LiGIoNs.

### 3.3. Method Evaluation

As mentioned above, LiGIoNs was evaluated using jackknife cross-validation. The workflow of the evaluation procedure is shown in **Figure 3**. The method’s ability to correctly identify the sequences is assessed each time and the procedure is repeated until all entries belonging to all subfamilies have been checked. The overall performance statistics of LiGIoNs are calculated afterwards.

**Figure 3.**
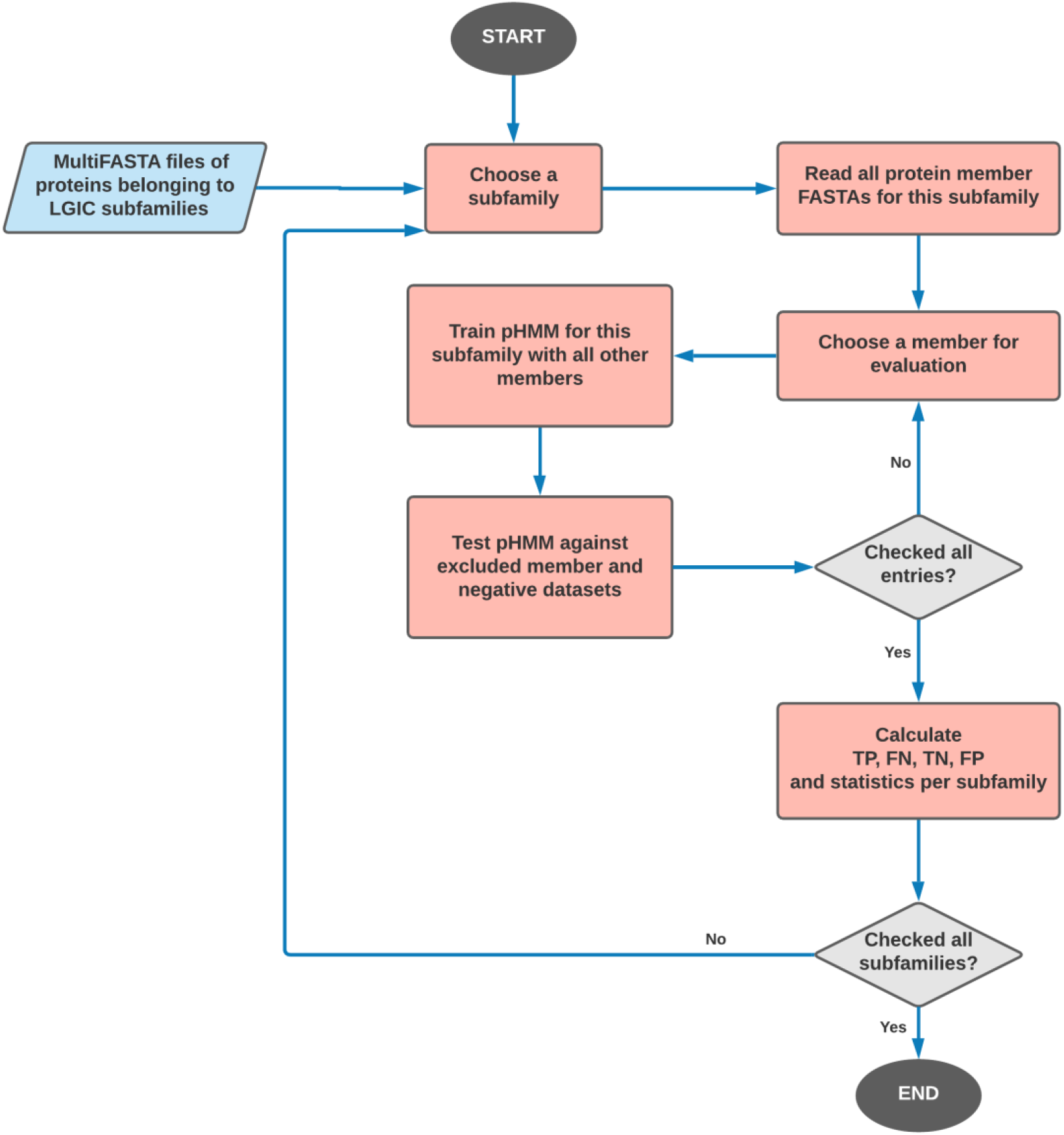
The workflow of the jackknife cross-validation experiment to assess the performance of LiGIoNs. Initially a subfamily of LGICs is chosen and all protein member FASTAs used for the training of the specific subfamily are selected. One protein sequence is used each time, which is left out, and the rest are used to create an MSA, train a pHMM and scan it against this one sequence, as well as, against the three negative datasets (**Supplementary Table 2**).

LiGIoNs performed very well during cross-validation, with high overall performance metrics against the globular and transmembrane negative test datasets (**Table 3** and **Table 4**). The results from the jackknife test showed the method’s ability to correctly identify pseudo-novel sequences as LGICs (Recall over 87% in most cases), except for the two smallest subfamilies (5-HT_3_ Receptors and ZACs). Additionally, no globular proteins and very few non-LGIC transmembrane proteins were erroneously classified as LGICs, with Precision being 100% and over 78% for each negative dataset respectively. Overall, the performance of LiGIoNs against the transmembrane negative dataset was good as the F-score was over 80% for most families, again with the two smallest subfamilies having a lower score but still over 72%. These values are indicative of the fact that pHMMs have a good ability to discriminate LGICs and non-LGICs, however they appear to underperform when classifying LGICs into the right subfamily (**Table 5**). For that reason, the method was also executed normally following the workflow described in **Section 3.1**, and proteins that have hits against multiple subfamilies were classified based on the highest overall score (see **Figure 1**). In that more realistic scenario, it became evident that LiGIoNs correctly classifies each known LGIC in the appropriate subfamily, as only a single LGIC classified in the wrong subfamily.

**Table 3.** Results from the cross-validation of LiGIoNs using the jackknife technique against the 953 globular proteins negative dataset

| *LGIC subfamily* | *Recall (%)* | *Precision (%)* | *F-score (%)* |
| --- | --- | --- | --- |
| Epithelial Sodium Channels (ENaCs) | 100.00 | 100.00 | 100.00 |
| P2X Receptors | 91.67 | 100.00 | 95.65 |
| Acid-sensing Ion Channels (ASICs) | 100.00 | 100.00 | 100.00 |
| Ionotropic Glutamate Receptors | 97.01 | 100.00 | 98.48 |
| Nicotinic Acetylcholine Receptors | 98.97 | 100.00 | 99.48 |
| 5-HT<sub>3</sub> Receptors | 60.00 | 100.00 | 75.00 |
| GABA<sub>A</sub> Receptors | 88.89 | 100.00 | 94.12 |
| Glycine Receptors | 88.24 | 100.00 | 93.75 |
| Zinc-activated Channels (ZACs) | 66.67 | 100.00 | 80.00 |
| IP3 Receptors | 87.50 | 100.00 | 93.33 |

**Table 4.** Results from the cross-validation of LiGIoNs using the jackknife technique against the 1508 transmembrane proteins of the negative dataset.

| *LGIC subfamily* | *Recall (%)* | *Precision (%)* | *F-score (%)* |
| --- | --- | --- | --- |
| Epithelial Sodium Channels (ENaCs) | 100.00 | 78.57 | 88.00 |
| P2X Receptors | 91.67 | 100.00 | 95.65 |
| Acid-sensing Ion Channels (ASICs) | 100.00 | 80.00 | 88.89 |
| Ionotropic Glutamate Receptors | 97.01 | 100.00 | 98.48 |
| Nicotinic Acetylcholine Receptors | 98.97 | 98.97 | 98.97 |
| 5-HT<sub>3</sub> Receptors | 60.00 | 100.00 | 75.00 |
| GABA<sub>A</sub> Receptors | 88.89 | 94.12 | 91.43 |
| Glycine Receptors | 88.24 | 100.00 | 93.75 |
| Zinc-activated Channels (ZACs) | 66.67 | 80.00 | 72.73 |
| IP<sub>3</sub> Receptors | 87.50 | 100.00 | 93.33 |

**Table 5.** Results from the cross-validation of LiGIoNs using the jackknife technique against LGICs belonging to other subfamilies than the one validated.

| *LGIC subfamily* | *Recall (%)* | *Precision (%)* | *F-score (%)* |
| --- | --- | --- | --- |
| Epithelial Sodium Channels (ENaCs) | 100.00 | 50.00 | 66.67 |
| P2X Receptors | 91.67 | 100.00 | 95.65 |
| Acid-sensing Ion Channels (ASICs) | 100.00 | 52.17 | 68.57 |
| Ionotropic Glutamate Receptors | 97.01 | 100.00 | 98.48 |
| Nicotinic Acetylcholine Receptors | 98.97 | 70.07 | 82.05 |
| 5-HT<sub>3</sub> Receptors | 60.00 | 5.17 | 9.52 |
| GABA<sub>A</sub> Receptors | 88.89 | 30.77 | 45.71 |
| Glycine Receptors | 88.24 | 45.45 | 60.00 |
| Zinc-activated Channels (ZACs) | 66.67 | 15.38 | 25.00 |
| IP<sub>3</sub> Receptors | 87.50 | 100.00 | 93.33 |

Even though most results presented above appear excellent, the pHMMs for 5-HT_3_ Receptors and ZACs perform moderately because the datasets for those LGIC sub-families are extremely small (**Supplementary Table 1**). An additional issue is that these subfamilies belong to the Cys-loop family of LGICs, so the ability to distinguish them from other members of the same family, primarily Nicotinic Acetylcholine Receptors, becomes inherently difficult (**Table 5**). The method’s ability to identify and characterize more proteins belonging to these subfamilies can be improved in the future, if LGICs belonging to proteomes that are evolutionary distant to mammals are annotated as such and are subsequently used during the creation of pHMMs. This is discussed further in Section 3.4.

LiGIoNs was also compared with IonChanPred 2.0 **[27]**. Based on the results presented in **Table 6**, it is obvious that our method outperforms IonChanPred 2.0 in the detection of LGICs. The positive dataset that was used to compare the two methods is the same as that presented in the original IonChanPred 2.0 publication **[27]**. For the negative set the original negative dataset presented in IonChanPred 2.0 was added to the two negative datasets presented earlier in this work (globular proteins and trans-membrane proteins other than LGICs). It should be noted that the two methods have different abilities, since, while IonChanPred 2.0 can detect LGICs, it lacks the ability to classify them into subfamilies. For this reason, the two methods are only compared for their ability to predict if a protein is an LGIC or not.

**Table 6.** Comparison of LiGIoNs with IonChanPred 2.0 in their ability to detect LGICs from their amino acid sequence.

| *Method* | *Recall (%)* | *Precision (%)* | *F-score (%)* |
| --- | --- | --- | --- |
| LiGIoNs | 93.33 | **97.22** | **95.24** |
| IonChanPred 2.0 | **100.00** | 40.43 | 57.58 |

This comparison shows that IonChanPred 2.0 has a recall of 100%, detecting all LGICs in the test set, but at the same time has very low precision, indicating that the high recall of the method is probably a result of overfitting to the positive set. In contrast, LiGIoNs can most often correctly identify LGICs, while not falsely detecting non-LGICs as LGICs. Overall, this demonstrates that LiGIoNs is a more balanced method, that avoids the issue of overfitting.

### 3.4. Application in reference proteomes

LiGIoNs was applied in all reference proteomes in UniProt for the detection of possible LGICs (**Supplementary Table 3**). Across the three superkingdoms, at least one potential LGIC was detected in almost half Eukaryotic proteomes, 304 Bacterial proteomes and only a single Archaeal proteome (**Table 7**). The FPR was calculated based on the transmembrane negative dataset and is 0.265%, which is higher than the percentage of LGICs found in Bacteria and Archaea, and with only 438 Eukaryotic proteomes surpassing this number. However, it should be noted that a) the true value of FPR is likely lower as transmembrane proteins are 30% or less of a whole proteome [40] and the FPR for globular proteins is 0%, and b) the threshold values for the pHMM are relatively high, possibly excluding more distant not yet annotated members.

**Table 7.** Results of the application of LiGIoNs in the reference proteomes of each superkingdom.

| *Superkingdom* | *Number of reference proteomes* | *Proteomes with at least one potential LGIC* | *Average number of LGICs* | *Average percentage of LGICs* | *Highest percentage of LGICs* |
| --- | --- | --- | --- | --- | --- |
| *Eukaryota* | 1586 | 726 | 87.23* | 0.33* | 1.09* |
| *Bacteria* | 7999 | 304 | 1.05 | 0.03 | 0.08 |
| *Archaea* | 330 | 1 | 1 | 0.04 | 0.04 |

\*excluding incomplete proteomes UP000265894 and UP000297026 with less than 500 proteins

Overall, the percentage of LGICs in Eukaryota is 0.33, but this number can vary significantly based on the kingdom and phylum. While LGICs are detected in all kingdoms and phyla, only some proteomes of Metazoan phyla have a higher percentage of LGICs than the calculated FPR. The results also showed that each LGIC subfamily varies in representation across the taxa, with Chordata being the only taxon with representatives of all subfamilies. 5-HT_3_ Receptors, important for serotonin neurotransmission, were found in Chordata and in Ecdysozoa, specifically in nematodes. ZACs, first discovered in humans and dogs [41], were found in Chordata including several mammals, bony fish and birds. Also, no ZACs were found in the rat and mouse proteomes, which is in accordance with the literature. Members of the subfamilies ASICs and ENaCs – both part of the ENaC/Degenerin family – were found in almost all Metazoan phyla, both vertebrate and invertebrate, as expected [42]. GABA_A_ receptors similarly were primarily limited in Metazoa – no members of this family were detected in plants despite the presence of GABA regulated ion channels in plants [43]. Additionally, 57 GABA_A_ receptors were predicted by LiGioNs in some bacterial proteomes. While this type of receptors has been found in bacteria [44], the percentage of predictions is less than the FPR. Next, it is known that plants have ionotropic glutamate receptors [45], and indeed most predicted LGICs in plants where assigned to the Ionotropic Glutamate Receptors subfamily. Glutamate receptors were also found in Metazoa and in several bacterial proteomes. Their presence in both eukaryotes and bacteria suggests that they emerged before the split from the common ancestor [46]. Another ancient subfamily is the IP3 Receptors that rose in eukaryotes before multicellularity according to previously published studies [47]. Possible LGICs belonging to this family were found in most eukaryotic kingdoms and phyla. Most predicted Nicotinic Acetylcholine Receptors belonged to Metazoa with fewer possible members found in other eukaryotic taxa and some bacteria. Even though it is thought that Acetylcholine has been synthesized since the beginning of life, receptors for this molecule have not been identified in any taxon apart from Metazoa [48]. Lastly, P2X Receptors were detected with LiGioNs, amongst other taxa, in Fungi [49] and Amoebozoa [50].

Approximately 61% of all detected LGICs are found in Chordata (253 proteomes) and all organisms with a percentage of LGICs higher than the FPR belonged to Metazoa. This is indicative of such organisms being better annotated and supports the notion that LiGIoNs could be overfitted to this type of data. Available evidence supports the existence of LGICs in several other taxa, such as bacteria [51,52], archaea [52] and fungi [49], however these receptors are not as of yet well studied and annotated, and are therefore underrepresented in protein sequence databases. LiGIoNs can easily be updated once new training data become available in order to improve its performance on evolutionary distant organisms. However, there is still significance in the results obtained from the current version of LiGioNs for organisms that are evolutionary distant to Metazoa. Specifically, despite the percentage of predicted prokaryotic LGICs not surpassing the FPR, a closer look at these 318 proteins (**Supplementary Table 3)** shows that the vast majority are potential LGICs or transporters, a closely related category of transmembrane proteins [53]. Many of them are also assigned to an LGIC-related Pfam entry (**Table 2**). Therefore, a more detailed examination of the results showed that they can serve as a steppingstone for further experimental validation of the function of prokaryotic proteins and could shed light in the evolution of this protein class of valuable pharmacological targets.

## 4. CONCLUSIONS

LiGIoNs is a fast and accurate method, which can detect Ligand-Gated Ion Channels from sequence alone and is therefore applicable to entire proteomes. LiGIoNs is the only publicly available method that classifies LGICs into one of the ten known subfamilies of these proteins. Moreover, LiGIoNs annotates predicted LGICs with information from Pfam, providing a full description of each sequence’s characteristics to the method’s users. LiGIoNs was applied to all UniProt reference proteomes. LGICs were detected in Eukaryota, Bacteria and Archaea, however the vast majority of receptors were found in Chordata.

In addition, the method we have developed is retrainable if more LGIC sequences become available in sequence databases. Retraining LiGIoNs when more LGICs are annotated, will allow us to increase the detection of LGICs in evolutionary distant proteomes than those used in the creation of the 10 core pHMMs. We plan to update the LGIClib pHMM library of LiGIoNs when new sequences are available, and this will allow us to build more descriptive profiles in the future and render the method more broadly applicable. Moreover, if new domains describing LGICs are added in Pfam, we plan to incorporate them in our method, as well. LiGIoNs is available at http://bioinformatics.biol.uoa.gr/ligions/.

## ABBREVIATIONS

*LGIC(s)*: Ligand-Gated Ion Channel(s)
*VGIC(s)*: Voltage-Gated Ion Channel(s)
*pHMM(s)*: profile Hidden Markov Models
*TM*: Transmembrane
*MSA*: Multiple Sequence Alignment
*MCC*: Matthew’s Correlation Coefficient

## ACKNOWLEDGEMENTS

The authors thank the National and Kapodistrian University of Athens for granting access to university premises and equipment.

## Author Contributions

Study design: KCN, ZIL, VAI; Conceptualization: AEA, KCN, GNP, ZIL; Method design and development: AEA, GNP, KCN; Web Application Design and Development: AEA, GNP; Web Application Quality Assurance: AEA, KCN, GNP, ZIL, VAI; Writing – original draft: AEA, KCN; Writing – review and editing: AEA, KCN, GNP, ZIL, VAI; Supervision: VAI.

